# Anisotropic regulation of lateral surface tension directs invagination in *C. elegans* gastrulation

**DOI:** 10.64898/2026.11.06.764797

**Authors:** Tokitaka Katayama, Kazunori Yamamoto, Toshikaze Chiba, Masayuki Imai, Fumio Motegi, Akatsuki Kimura

**Affiliations:** The Graduate University for Advanced Studies, Japan; National Institute of Genetics, Japan; Institute for Genetic Medicine, Hokkaido University, Japan; Department of Physics, Graduate School of Science, Tohoku University, Japan; Department of Theoretical Physics, Institut Jožef Stefan, Slovenia

## Abstract

Epithelial invagination is commonly attributed to actomyosin-driven apical constriction, but how the direction of tissue deformation is determined remains unclear. Here, we show that gastrulation in *Caenorhabditis elegans* involves anisotropic invagination, with preferential anterior–posterior convergence of neighboring cells. Laser ablation revealed no detectable directional bias in apical cortical recoil, suggesting that anisotropy in apical cortical tension is unlikely to be the primary determinant of the observed directional deformation. Instead, HMR-1/E-cadherin exhibited anisotropic accumulation preferentially at specific lateral cell–cell interfaces, where its enrichment is associated with junctional geometry consistent with reduced interfacial surface tension. Three-dimensional mechanical simulations showed that selectively reducing tension at these interfaces reproduces the observed anisotropic deformation and markedly enhances invagination, whereas uniform tension reduction does not. These findings reveal that spatially patterned lateral surface tension cooperates with apical constriction to control the direction and extent of invagination, uncovering a mechanical role for cell adhesion beyond force transmission.

## Introduction

Multicellular animals form three-dimensional, tube-like structures such as the trachea, stomach, and gut by folding two-dimensional, sheet-like tissues^1–3^. Invagination is a fundamental morphogenetic process through which epithelial sheets fold to generate three-dimensional structures during embryogenesis and tissue formation^4^. In many systems, including gastrulation and neurulation, invagination has been primarily explained by apical constriction driven by actomyosin contractility^5–7^. In this process, apically localized non-muscle myosin II contracts the cortical actin network beneath the plasma membrane, thereby reducing the apical surface and generating epithelial bending^8,9^. This framework has provided a successful mechanical description of epithelial folding across diverse organisms^5–7^.

Invagination occurs in at least two geometrically distinct forms. In one form, invagination extends along an elongated region of the epithelium, as exemplified by *Drosophila* ventral furrow formation^10–12^ and vertebrate neural tube formation^3^ (hereafter referred to as “furrow-like invagination”). In the other, invagination occurs within a spatially localized region of the epithelium, as exemplified by sea urchin gastrulation^13–16^ (hereafter referred to as “punctate invagination”). In the present study, we focus on punctate invagination. Punctate invagination is often described as an approximately axisymmetric process around the apico**–**basal axis passing through the invagination site (hereafter, the invagination axis). Indeed, approximately axisymmetric tissue deformation around the invagination axis has been observed during sea urchin gastrulation^16,17^. Accordingly, mechanical analyses of punctate invagination have generally focused on cross-sectional planes containing the invagination axis, an approach that is particularly suitable for describing axisymmetric deformation. However, whether punctate invagination necessarily proceeds symmetrically within the epithelial plane, which is approximately perpendicular to the invagination axis, remains largely unexplored.

*C. elegans* gastrulation provides an attractive system for investigating the mechanics of punctate invagination for two reasons: its well-characterized molecular mechanisms and the distinctive geometry of the embryo. Gastrulation begins at the 26-cell stage with the internalization of two endodermal precursor cells, Ea and Ep, from the ventral surface toward the interior of the embryo^18–20^. During this process, apical surfaces of Ea and Ep constrict, accompanied by the convergence of surrounding cells toward the invaginating site. This apical constriction is driven by NMY-2, the *C. elegans* non-muscle myosin II, which accumulates at the apical surfaces of Ea and Ep and generates contractile forces^19,21^. Contractile movements of the apical actomyosin network occur before the onset of apical constriction, but apical constriction begins only when these movements become mechanically coupled to the cell boundaries. This mechanism, known as the molecular clutch model, explains how actomyosin contractility is converted into apical surface constriction^21^. In addition to contractile proteins, cell adhesion molecules, including the E-cadherin homologue HMR-1 and the L1CAM homologue SAX-7, are required for successful gastrulation^22^. By mechanically linking neighboring cells, these cell adhesion molecules may facilitate the transmission of forces generated by Ea/Ep apical constriction to surrounding cells, contributing to their convergence toward the invagination site^21,23^. Importantly, unlike the idealized axisymmetric invagination models of punctate invagination, the geometry of the *C. elegans* embryo is inherently anisotropic. The embryo is already elongated along the anterior**–**posterior axis, and the two internalizing cells are arranged along this axis, thereby defining distinct anterior–posterior (AP) and left–right (LR) directions before invagination begins^19,24,25^. These geometric features raise the possibility that punctate invagination may not proceed symmetrically within the epithelial plane.

Here, we first asked whether punctate invagination during *C. elegans* gastrulation indeed proceeds symmetrically within the epithelial plane. We sought to determine, if not, how such anisotropic deformation is mechanically generated and whether it actively contributes to tissue folding.

## Results

### Ea/Ep internalization is accompanied by preferential anterior–posterior, rather than left-right, convergence of neighboring cells around the invaginating cells

To determine whether punctate invagination proceeds symmetrically within the epithelial plane, we first analyzed tissue deformation surrounding the invaginating Ea and Ep cells. We visualized cell movements in living embryos expressing PH::mCherry, which labels the plasma membrane, by confocal microscopy (Figures 1A and 1B). Throughout this study, we defined time 0 as the birth of the MS granddaughter cells (MSaa, MSap, MSpa, and MSpp), corresponding to the onset of invagination. The completion of invagination was defined as the time when the Ea and Ep cells entered mitosis.

Embryos mounted before the four-cell stage reproducibly adopted an orientation that provided a lateral view during gastrulation (Figure 1A; Video S1). At the four-cell stage, the four blastomeres become arranged within the dorsal–ventral plane, resulting in a flattened embryo whose shortest dimension is along the LR axis^25^. Consequently, the embryo preferentially settles with the LR axis perpendicular to the coverslip. This orientation was favored even when embryos were mounted using 30 µm spacer beads to minimize compression. However, the lateral view is not well suited for analyzing deformation of the apical surfaces of Ea and Ep cells during their internalization.

To directly visualize the tissue deformation within the epithelial plane, we developed a mounting procedure in which embryos were dissected from adult hermaphrodites, allowed to develop to the four-cell stage, and then manually oriented with the ventral side facing the objective before mounting. This procedure enabled reproducible imaging from the ventral side (Figure 1B; Video S2). Using these image stacks, cell boundaries were segmented based on the membrane fluorescence signal (the plasma membrane marker PH::mCherry), and three-dimensional cell geometries were reconstructed. Individual cells were identified manually, and centroid coordinates were calculated for subsequent quantitative analyses (Figure 1B’).

To investigate whether tissue deformation is axisymmetric around the invagination axis or exhibits anisotropy within the epithelial plane, we quantified changes in the positions of the six cells surrounding Ea and Ep, comparing deformation along the AP and LR directions. For each of the 15 possible pairs of the six surrounding cells, we calculated the vector connecting their three-dimensional centroids and determined its AP and LR components to quantify the intercellular distance along each axis (Figure 1C). If tissue deformation were isotropic within the epithelial plane, comparable shortening would be expected along the AP and LR axes. However, cell–cell distances decreased predominantly along the AP axis, with reductions reaching up to 15 µm, whereas only limited shortening (<5 µm) was observed along the LR axis. These measurements indicate that Ea/Ep internalization is accompanied by preferential AP convergence of neighboring cells. Thus, punctate invagination in *C. elegans* deviates from axisymmetric deformation around the invagination axis, revealing an unexpected anisotropy in tissue deformation within the epithelial plane during gastrulation. Within the framework of apical constriction-driven invagination, this preferential AP convergence suggests that the apical surfaces of Ea and Ep undergo greater shortening along the AP axis than along the LR axis. This raises the question of how such anisotropic deformation is mechanically generated.

**Figure 1.**
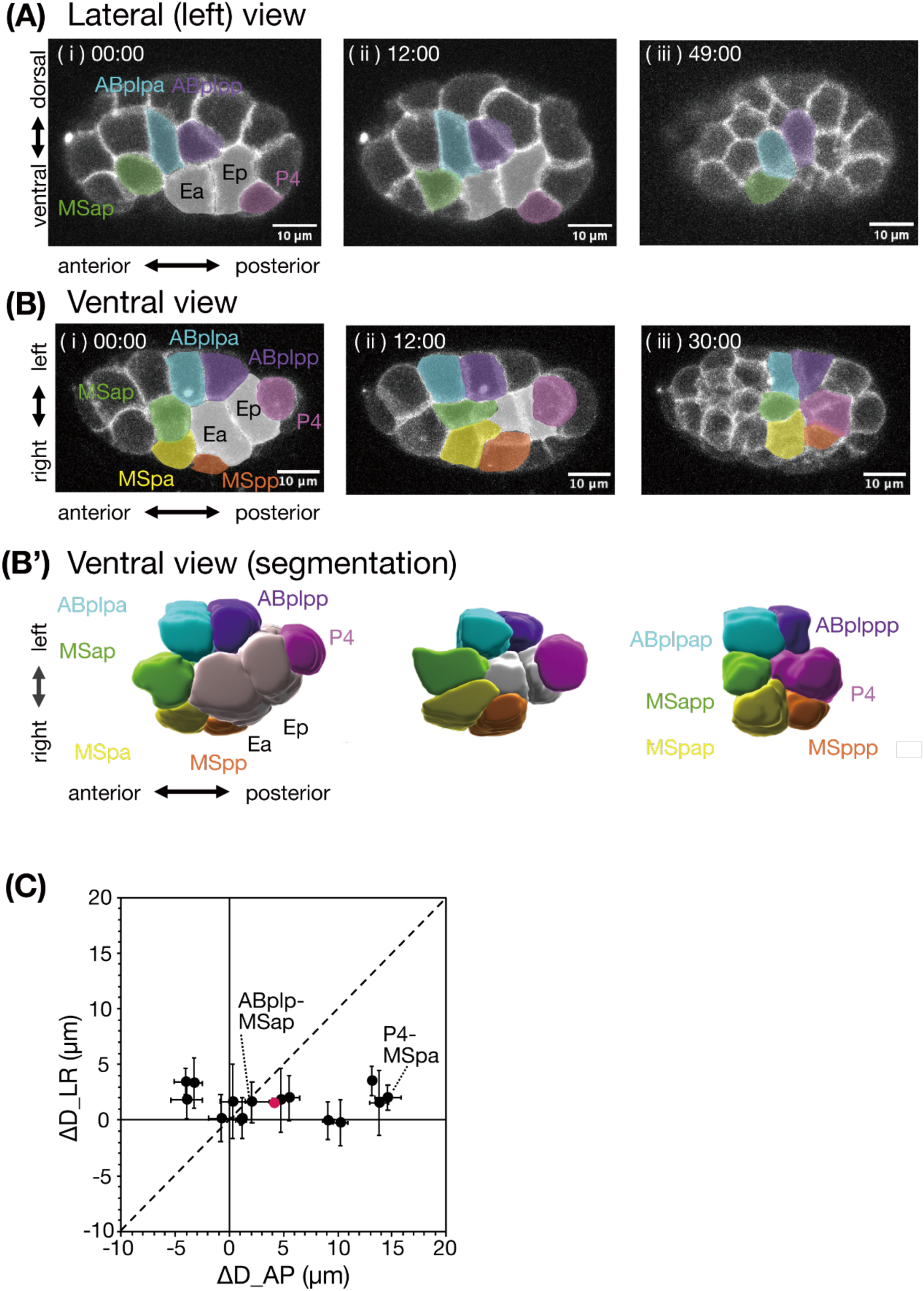
Ea/Ep internalization is accompanied by preferential anterior–posterior convergence of neighboring cells. **(A)** Lateral (left-side) view of *C. elegans* gastrulation. Panels (i–iii) show representative images at the onset of Ea/Ep internalization (time 0; immediately after division of the MS daughter cells), during internalization, and upon completion of internalization (immediately before Ea/Ep division), respectively. Time stamps are shown in minutes:seconds (min:s). **(B)** Ventral view of gastrulation. Panels (i–iii) correspond to the same developmental stages shown in (A). The embryo shown in (B) is different from that in (A). **(B′)** Three-dimensional reconstruction of the embryo shown in (B). Cell boundaries were segmented based on cell membrane (PH::mCherry) fluorescence, and individual blastomeres were identified manually. Panels (i–iii) correspond to the same time points as in (B). **(C)** Quantification of the relative movements of the six cells surrounding the internalizing Ea and Ep cells. These surrounding cells were defined as the six cells that overlie the apical surfaces of Ea and Ep at the completion of internalization, as viewed from the ventral side. For each embryo, changes in the relative positions of all 15 possible pairs of the six surrounding cells were quantified separately along the AP and LR axes. ΔD_AP and ΔD_LR represent the reductions in the AP and LR components of the intercellular distances, respectively, from the onset to the completion of internalization. Positive values indicate shortening, whereas negative values indicate separation along the corresponding axis. The dashed diagonal line indicates ΔD_AP = ΔD_LR. Points below the diagonal line indicate greater shortening along the AP axis than along the LR axis. Black circles indicate the mean values for individual cell pairs (n = 5 embryos), with error bars representing standard deviation. The P4–MSpa and ABplp–MSap pairs are highlighted as examples showing pronounced and limited AP shortening, respectively. The magenta circle indicates the overall mean of all 75 measurements (15 cell pairs per embryo, n = 5 embryos).

### Video S1. Ea/Ep internalization viewed from the lateral side

Time-lapse fluorescence imaging of Ea/Ep internalization during *C. elegans* gastrulation, viewed from the lateral (left) side. Cell membranes are visualized using PH::mCherry. Images are maximum-intensity projections of Z-stacks. Time is indicated in minutes relative to the birth of MS daughter cells. Scale bar, 10 µm.

### Video S2. Ea/Ep internalization viewed from the ventral side

Time-lapse fluorescence imaging of Ea/Ep internalization during *C. elegans* gastrulation, viewed from the ventral side. Cell membranes are visualized using PH::mCherry. Images are maximum-intensity projections of Z-stacks. Time is indicated in minutes relative to the birth of MS daughter cells. Scale bar, 10 µm.

### Apical cortical recoil shows no detectable anisotropy along the AP and LR axes

The preferential AP convergence of surrounding cells observed in Figure 1 suggests anisotropic shortening of the apical surfaces during invagination. One possible explanation for this anisotropic deformation is that apical cortical tension is stronger along the AP axis than along the LR axis. To test this possibility, we examined the directional dependence of cortical recoil on the apical surface of Ep cells using laser ablation. Previous studies have used laser line ablation to compare cortical tension along the AP axis and its orthogonal axis in one-cell-stage *C. elegans* embryos, with the two directions measured in separate embryos^26,27^. To avoid comparisons between embryos and accommodate the small apical surface of Ep, we adapted this approach by using point ablation to assess the directional bias of cortical recoil along the AP and LR axes from a single ablation event (Figure 2A). Laser ablation locally releases cortical tension, causing the surrounding cortex to recoil away from the ablation site. Assuming comparable mechanical responses to tension release along the AP and LR axes, greater recoil along one axis would suggest stronger pre-existing cortical tension along that axis.

**Figure 2.**
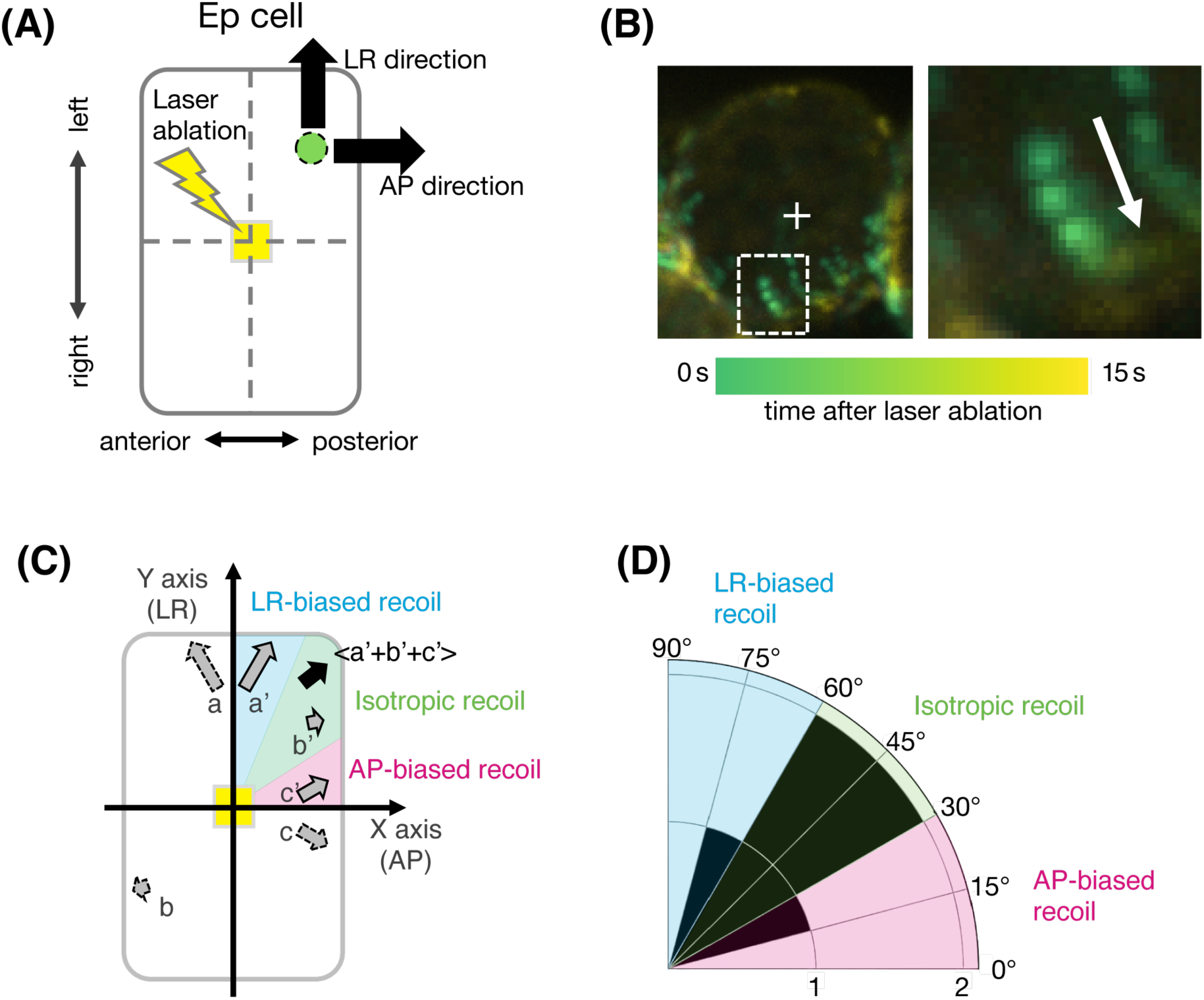
Laser ablation reveals no detectable directional bias in apical cortical recoil. **(A)** Schematic of the laser ablation assay used to evaluate cortical tension on the apical surface of the Ep cell. A single point on the apical cortex was irradiated with a laser (yellow rectangle), and the recoil of cortical NMY-2 puncta surrounding the ablation site was analyzed to determine whether cortical tension is preferentially oriented along the AP or LR axis. **(B)** Representative ventral view of an Ep cell expressing NMY-2::mBaoJin before and after laser ablation. Consecutive time-lapse images acquired at 1-s intervals were pseudo-colored according to acquisition time and overlaid into a single image. The enlarged view (right) shows representative displacement of a cortical NMY-2 punctum away from the ablation site (arrow). **(C)** Quantification of recoil direction. NMY-2 puncta were tracked between consecutive frames using a particle-tracking algorithm. Displacement vectors were expressed relative to the ablation site, with the AP axis defined as the x-axis and the LR axis as the y-axis. To compare directional bias independently of the original quadrant, vectors were reflected into the first quadrant while preserving their orientation relative to the AP and LR axes. The mean displacement vector was then calculated. The angle of the mean displacement vector represents the preferred recoil direction (0°, AP; 90°, LR; 45°, no directional bias), whereas its magnitude represents the extent of cortical recoil. **(D)** Distribution of the representative recoil angles (stars in Figure S1). Recoil angles were centered around 45°, suggesting no preferential recoil along either the AP or LR axis.

**Figure S1.**
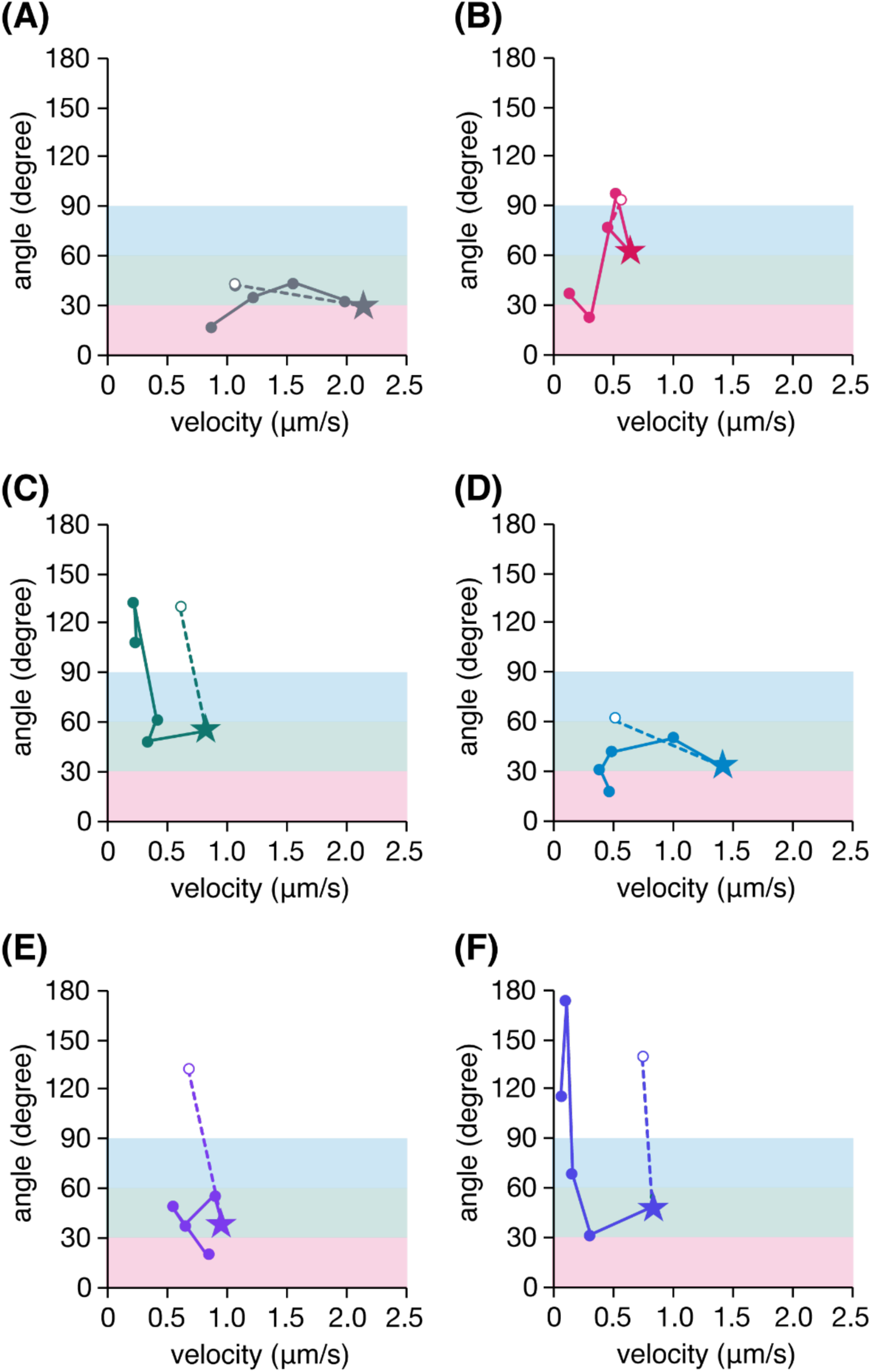
Temporal changes in the direction and magnitude of the mean displacement vector of NMY-2 puncta following laser ablation. **(A–F)** Each panel shows data from a different embryo (n = 6 embryos in total). The horizontal axis shows the magnitude of the mean displacement vector divided by the time interval between consecutive frames (1 s), expressed as velocity (μm/s). The mean displacement vector was calculated by vectorially averaging the displacements of NMY-2 puncta, as described in the main text (Figure 2C). The vertical axis shows the angle of the mean displacement vector relative to the AP axis. Each point represents a measurement at a single time point. Open circles indicate measurements immediately before laser ablation, whereas filled circles indicate measurements at four consecutive time points after ablation, at 1-s intervals. Dashed lines connect the pre- and post-ablation measurements. Stars indicate the time points at which the velocity reached its maximum in each embryo. The corresponding angles were used as the representative recoil angles in Figure 2D.

Embryos were mounted in the ventral-side orientation described in Figure 1B, and laser ablation was performed 22 min after Ea/Ep birth, corresponding to the time of the most rapid decrease in apical surface area on average across embryos. Cortical NMY-2 puncta moved away from the ablation site immediately after irradiation (Figure 2B). To quantify the recoil direction, NMY-2 puncta were tracked between consecutive image frames using a particle-tracking algorithm (Figure 2C). Displacement vectors were expressed in a coordinate system centered on the ablation site, with the AP axis defined as the x-axis and the LR axis as the y-axis. To compare directional bias independently of the quadrant in which puncta were located, vectors from all four quadrants were reflected into the first quadrant while preserving their orientation relative to the AP and LR axes. The mean displacement vector was then calculated by summing all transformed vectors and dividing by the number of vectors. The angle of the mean displacement vector indicates the preferred recoil direction. Angles between 0° and 90° indicate net movement away from the ablation site, with 0° corresponding to the AP axis, 90° to the LR axis, and 45° indicating no directional bias between the two axes. The magnitude of the mean displacement vector, expressed as velocity (μm/s) because the frame interval was 1 s, reflects both the magnitude of individual punctum displacements and their directional alignment, as displacements in different directions tend to cancel each other when averaged as vectors.

Before laser ablation, the magnitude of the mean displacement vector was small, corresponding to a low velocity, and fluctuated randomly in direction (Figure S1). Immediately after ablation, the velocity of the mean displacement vector increased sharply, suggesting that individual NMY-2 puncta underwent larger, more directionally aligned displacements. The velocity subsequently declined. Because the timing of maximal recoil varied slightly among embryos, we selected the frame with the highest velocity of the mean displacement vector for each embryo (Figure S1, star) and used the corresponding angle of the mean displacement vector as the representative recoil angle (Figure 2D). At this time point, the angle was between 0° and 90°, indicating a net outward movement of NMY-2 puncta away from the ablation site, consistent with rapid cortical recoil. Recoil angles clustered around 45°, indicating that cortical recoil occurred without a preferential orientation toward either the AP or LR axis. These results are inconsistent with the above hypothesis in which anisotropic deformation arises from stronger cortical tension along the AP axis. Instead, these findings suggest that anisotropy in apical cortical tension is unlikely to be the primary determinant of the observed directional deformation, pointing to the involvement of another mechanism during gastrulation.

### E-Cadherin is enriched at specific lateral interfaces, suggesting spatial differences in surface tension *in vivo*

Having found no detectable anisotropy in apical cortical recoil (Figure 2), we next considered whether differences in the mechanical properties of lateral cell–cell interfaces contribute to anisotropic deformation during gastrulation. Previous studies have shown that E-cadherin-mediated cell adhesion is required for *C. elegans* gastrulation^22^. E**-**cadherin-mediated adhesion has two distinct mechanical roles. One is to mechanically couple neighboring cells, thereby transmitting contractile forces across cell–cell junctions^28^. This role has been considered the primary contribution of E-cadherin to *C. elegans* gastrulation^21,22^. The other is to reduce surface tension at cell–cell interfaces^29,30^. Surface tension at these interfaces reflects the balance between cortical contractility, which tends to reduce the contact area, and cell–cell adhesion, which favors its expansion. Although this second role is well established, its contribution to gastrulation remains unclear. We therefore asked whether E-cadherin-mediated reduction of lateral interfacial tension contributes to the anisotropic deformation observed during gastrulation.

We first examined the localization of HMR-1, the *C. elegans* E-cadherin, at lateral cell–cell interfaces around Ea and Ep using embryos expressing HMR-1::GFP. Strikingly, ventral-view imaging revealed that HMR-1 was not uniformly distributed among these interfaces but was preferentially enriched at specific cell–cell contacts (Figure 3A).

**Figure 3.**
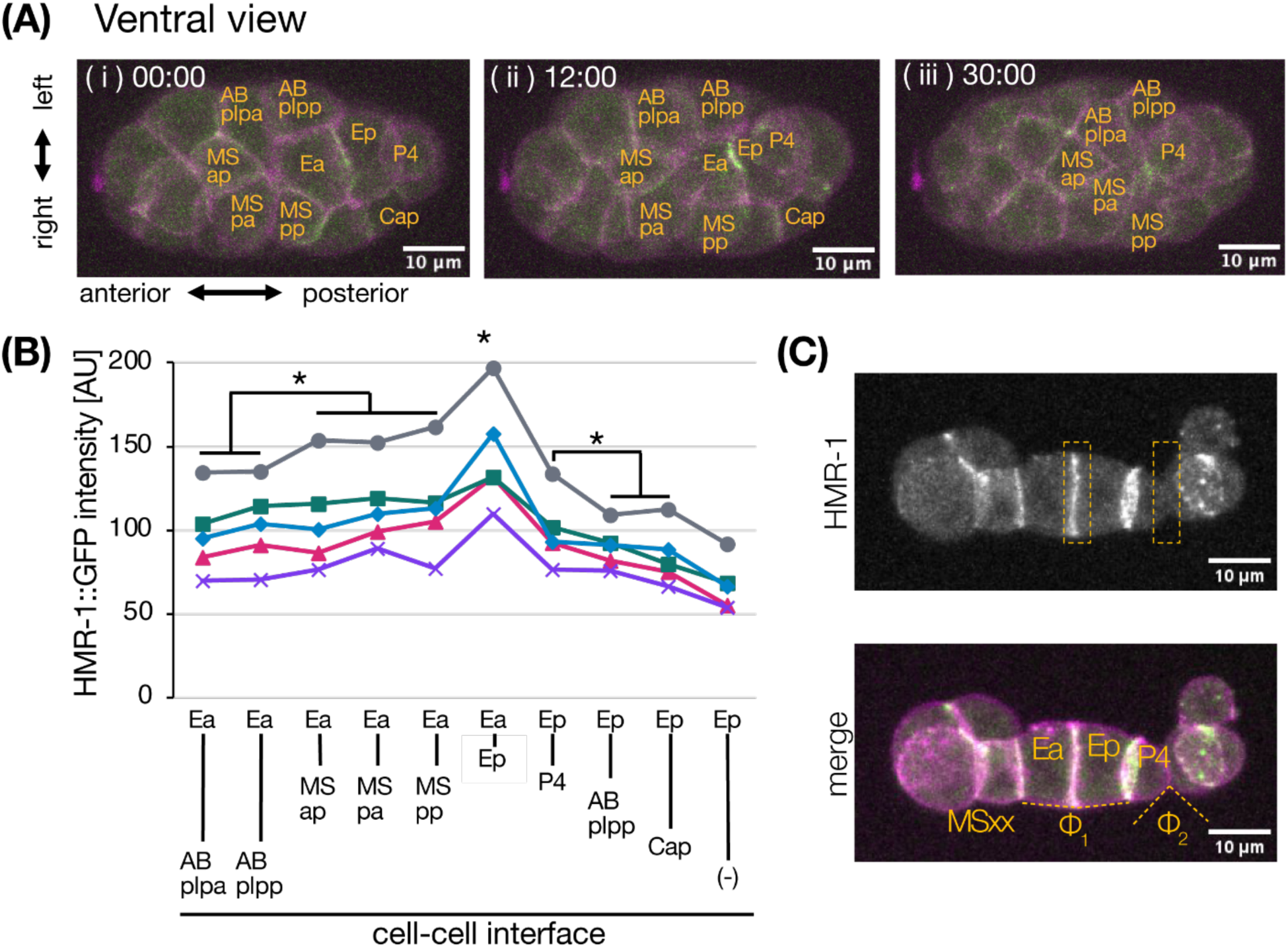
HMR-1 is preferentially localized to specific lateral interfaces, which exhibit junctional geometry consistent with reduced surface tension. **(A)** Representative ventral view of an intact embryo at the 26-cell stage expressing HMR-1::GFP (green) and PH::mCherry (cell membrane marker, magenta). Panels (i–iii) correspond to the same developmental stages shown in panels (i–iii) of Figure 1A. **(B)** Quantification of HMR-1::GFP intensity at lateral cell–cell interfaces. Each value represents the mean HMR-1::GFP fluorescence intensity at the indicated cell surface, averaged over the period from the onset to the completion of Ea/Ep internalization. The two cells forming each interface are indicated below the x-axis. Ep–(−) denotes the non-contacting lateral surface of Ep. Each color–symbol combination represents an individual embryo (n = 5 embryos), and measurements from the same embryo are connected by lines. The Ea–Ep interface exhibited the highest HMR-1::GFP fluorescence intensity in all five embryos, a pattern that was statistically significant under the null hypothesis of random ranking. Brackets indicate two additional group comparisons: (i) the Ea–MS interfaces (Ea–MSap, Ea–MSpa, and Ea–MSpp) versus the Ea–AB interfaces (Ea–ABplpa and Ea–ABplpp), and (ii) Ep–P4 versus Ep–ABplpp and Ep–Cap interfaces. Statistical significance was assessed using the respective tests described in the Methods. *P < 0.05. **(C)** Comparison of junctional angles at cell–cell interfaces with different levels of HMR-1 enrichment. Embryos were separated into AB and P1 blastomeres at the two-cell stage following eggshell removal. Isolated P1 blastomeres were cultured to generate descendant cells, including MSxx, Ea, Ep, and P4. The upper grayscale image shows HMR-1::GFP fluorescence, whereas the lower merged image shows HMR-1::GFP in green and the cell membrane marker PH::mCherry in magenta. Orange dashed rectangles highlight regions with different levels of HMR-1 enrichment. Junction angles were compared at HMR-1-enriched interfaces (e.g., Ea–Ep, Φ₁ = 196°) and interfaces with lower HMR-1 enrichment (e.g., Φ₂ = 100°). The markedly larger junctional angles at HMR-1-enriched interfaces are consistent with lower interfacial tension, assuming comparable mechanical properties of the surrounding cell surfaces.

To characterize this spatially restricted localization quantitatively, we measured the mean fluorescence intensity of HMR-1::GFP at individual lateral cell–cell interfaces extending perpendicular to the apical surfaces of Ea and Ep (Figure 3B). Ea contacts six neighboring cells: Ep, three MS-lineage cells (MSap, MSpa, and MSpp), and two AB-lineage cells (ABplpa and ABplpp). Ep contacts four neighboring cells (Ea, P4, ABplpp, and Cap) and also has a lateral surface that is not in contact with another cell (indicated as (–) in Figure 3B). Among these interfaces, the Ea–Ep interface exhibited the highest HMR-1::GFP fluorescence intensity (Figure 3B). At the interfaces of Ea, HMR-1::GFP fluorescence was consistently and significantly higher at contacts with MS-lineage cells than at those with AB-lineage cells. At the interfaces of Ep, HMR-1::GFP fluorescence was higher at the Ep–P4 interface than at the interfaces with ABplpp or Cap (Figure 3B). Together, these results demonstrate that HMR-1 is preferentially enriched at specific lateral cell–cell interfaces, particularly those between MS-lineage cells and Ea, between Ea and Ep, and between Ep and P4.

We next asked whether cell–cell interfaces with higher HMR-1 enrichment exhibit lower interfacial surface tension *in vivo*, as expected from stronger cell–cell adhesion. Junctional angles provide a geometric indication of relative interfacial tension, with larger angles indicating lower tension at the corresponding cell–cell interface, assuming comparable mechanical properties of the surrounding cell surfaces and other factors, such as intracellular pressure^23,29^. We therefore examined junctional geometry in isolated P1 blastomeres cultured to generate their descendants, including MSxx, Ea, Ep, and P4 (Figure 3C). The Ea–Ep interface, which showed strong HMR-1 enrichment in both intact embryos (Figures 3A and 3B) and isolated blastomeres (orange dashed rectangles in Figure 3C), exhibited a markedly larger junctional angle (Φ₁ in Figure 3C) than an interface with lower HMR-1 enrichment (Φ₂ in Figure 3C). This qualitative observation is consistent with the idea that interfacial surface tension is lower at HMR-1-enriched cell–cell interfaces.

### A three-dimensional mechanical model separates cell–cell connectivity from interfacial surface tension

Having found that HMR-1-enriched lateral interfaces exhibit junctional geometries consistent with reduced surface tension *in vivo* (Figure 3), we next asked whether spatial anisotropy in lateral surface tension could account for the anisotropic deformation observed during gastrulation. To address this question, we constructed a three-dimensional mechanical model using Surface Evolver^31^, which simulates shape changes by minimizing surface energy under prescribed mechanical constraints, including surface tension and volume conservation. Surface Evolver has been widely used to investigate the mechanics of tissue morphogenesis^32–35^.

A key feature of our model is that cell–cell connectivity and interfacial surface tension can be controlled independently. Rather than representing the interface between neighboring cells as a single shared surface, we modeled two contacting surfaces separately and connected adjacent cells using multiple short elastic elements. These elements maintain mechanical coupling between neighboring cells, representing one role of E-cadherin-mediated adhesion, while surface tension can be independently assigned to each contact surface to represent the other role.

Before applying this framework to gastrulation, we examined its mechanical behavior using a simple two-cell system. We first simulated two cells sharing a single contact surface, in which the equilibrium junction angle is predicted by the Young–Dupré relation^29,36^. As expected, decreasing the relative surface tension at the contact surface increased the junction angle, whereas increasing it decreased the angle (**Figures S2A–S2C**).

We then examined the alternative representation used in our model, in which neighboring cells have independent contact surfaces connected by short elastic elements. Although this representation produced smaller junction angles than the shared-surface model at the same relative surface tension, it preserved the same qualitative dependence on surface tension: increasing the tension at the contact surfaces decreased the junctional angle (**Figures S2D–S2G**). Thus, the model reproduced the expected qualitative relationship between interfacial surface tension and junction angle while allowing cell–cell connectivity and surface tension to be manipulated independently. We therefore adopted this framework for the subsequent gastrulation simulations.

**Figure S2.**
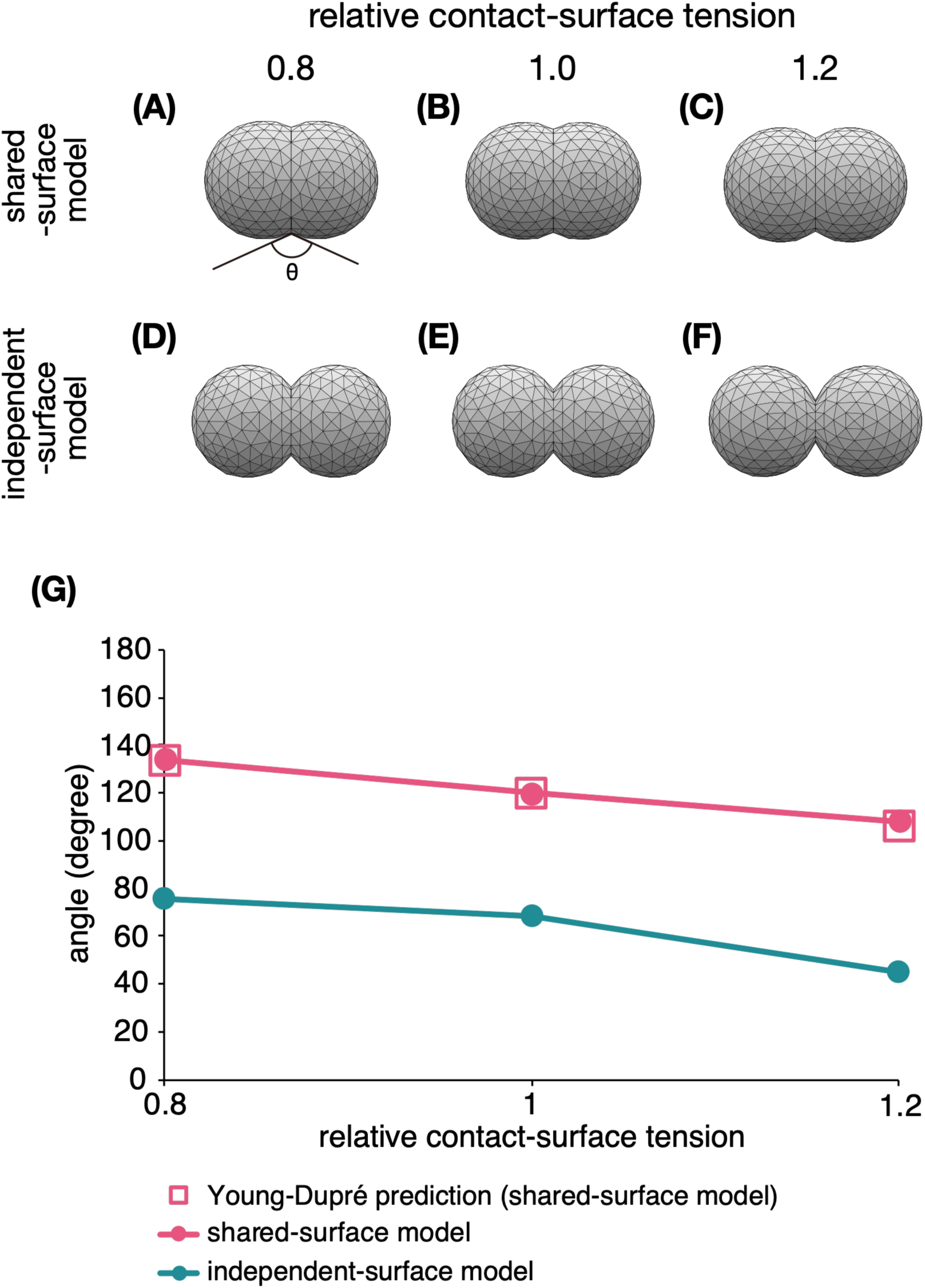
Dependence of junctional angles on contact-surface tension in two-cell simulations using Surface Evolver. **(A–C)** Two-cell simulations in which neighboring cells share a single contact surface. The relative surface tension at the contact surface was set to 0.8 (A), 1.0 (B), or 1.2 (C), while the tension at non-contact surfaces was maintained at 1.0. The equilibrium junction angle (θ) increased as the relative surface tension at the contact surface decreased, consistent with the Young–Dupré relation. **(D–F)** Two-cell simulations in which neighboring cells have independent contact surfaces connected by short elastic elements. The relative surface tension at the contact surfaces was set to 0.8 (D), 1.0 (E), or 1.2 (F), while the tension at non-contact surfaces was maintained at 1.0. Although the junction angles differed from those in the shared-surface model, they showed the same qualitative dependence on contact-surface tension. **(G)** Equilibrium junction angles plotted against relative surface tension at the contact surfaces. Pink symbols and lines indicate the shared-surface model, and teal symbols and lines indicate the independent-surface model with elastic connections. Open squares indicate the theoretical predictions based on the Young–Dupré relation. The shared-surface model closely reproduced the theoretical predictions. The independent-surface model also reproduced the expected qualitative relationship between contact-surface tension and junction angle, with junction angles decreasing as contact-surface tension increased.

### Anisotropic reduction of lateral surface tension generates anisotropic apical deformation

Having found that specific lateral cell–cell interfaces are associated with reduced surface tension *in vivo* (Figure 3), we next asked whether this anisotropic reduction in surface tension could generate anisotropic apical deformation consistent with the preferential AP convergence observed during gastrulation (Figure 1C). To capture the essential mechanics of gastrulation, we constructed a minimal four-cell model consisting of Ea, Ep, MSap, and P4, which constitute the core cellular configuration during *C. elegans* gastrulation^18,37^. Connectivity was maintained by elastic elements, whereas surface tension could be independently assigned to different cell surfaces.

The simulation was initialized with four connected cubic cells. When all surfaces were assigned a uniform baseline surface tension of 1.0, energy minimization caused the cells to relax into a configuration of approximately spherical cells, representing the pre-invagination state. To define the mechanical properties of the model, we classified the 24 surfaces of the four cells into four categories: the apical surfaces of Ea and Ep, lateral contact surfaces associated with high HMR-1 enrichment, lateral contact surfaces associated with low HMR-1 enrichment, and the remaining (“other”) surfaces (Figure 4A). Surface tension was then assigned to the individual surfaces according to the conditions specified for each model (Table 1). To simulate apical constriction, we progressively increased the surface tension of the apical surfaces of the Ea and Ep from 1.0 to 2.0, based on the established role of actomyosin contractility in driving apical constriction during gastrulation.

We first examined a ‘selective relaxation model,’ in which surface tension was reduced to 0.8 at the three lateral cell–cell interfaces associated with HMR-1 enrichment (MSap–Ea, Ea–Ep, and Ep–P4), while the tension of all other non-apical surfaces was maintained at 1.0 (Figures 4B, 4C, and Table 1). As apical surface tension increased, the apical surfaces of Ea and Ep underwent anisotropic deformation, with preferential shortening along the AP axis and relative elongation along the LR axis (Figure 4D; Video S3). This deformation was consistent with the preferential AP convergence of surrounding cells observed *in vivo* (Figure 1).

To determine whether this anisotropic deformation specifically requires a spatially patterned reduction in lateral surface tension, we compared the selective relaxation model with two control models in which lateral surface tension was uniform across Ea and Ep. In the ‘no-relaxation model,’ all non-apical surfaces retained the baseline tension of 1.0 (Figures 4E, 4F and Table 1). In the ‘uniform relaxation model,’ surface tension was reduced to 0.8 across all lateral surfaces of Ea and Ep, including those exposed to the exterior, while the tension of all remaining non-apical surfaces was maintained at 1.0 (Figures 4H, 4I and Table 1). In both models, apical surface tension was increased from 1.0 to 2.0, as in the selective relaxation model. Unlike the selective relaxation model, neither control model reproduced the preferential AP shortening observed in the selective relaxation model (Figures 4G and 4J; Videos S4 and S5).

To quantify the directional deformation of the simulated apical surfaces, we measured changes in the distances between opposing boundaries of the Ea and Ep apical surfaces along the AP and LR axes (Figure 4K). The selective relaxation model showed preferential shortening along the AP axis, whereas the no-relaxation and uniform relaxation models exhibited little directional bias. These results demonstrate that spatially patterned reduction of lateral surface tension can generate anisotropic apical deformation consistent with the preferential AP convergence observed *in vivo* (Figure 1C).

**Figure 4.**
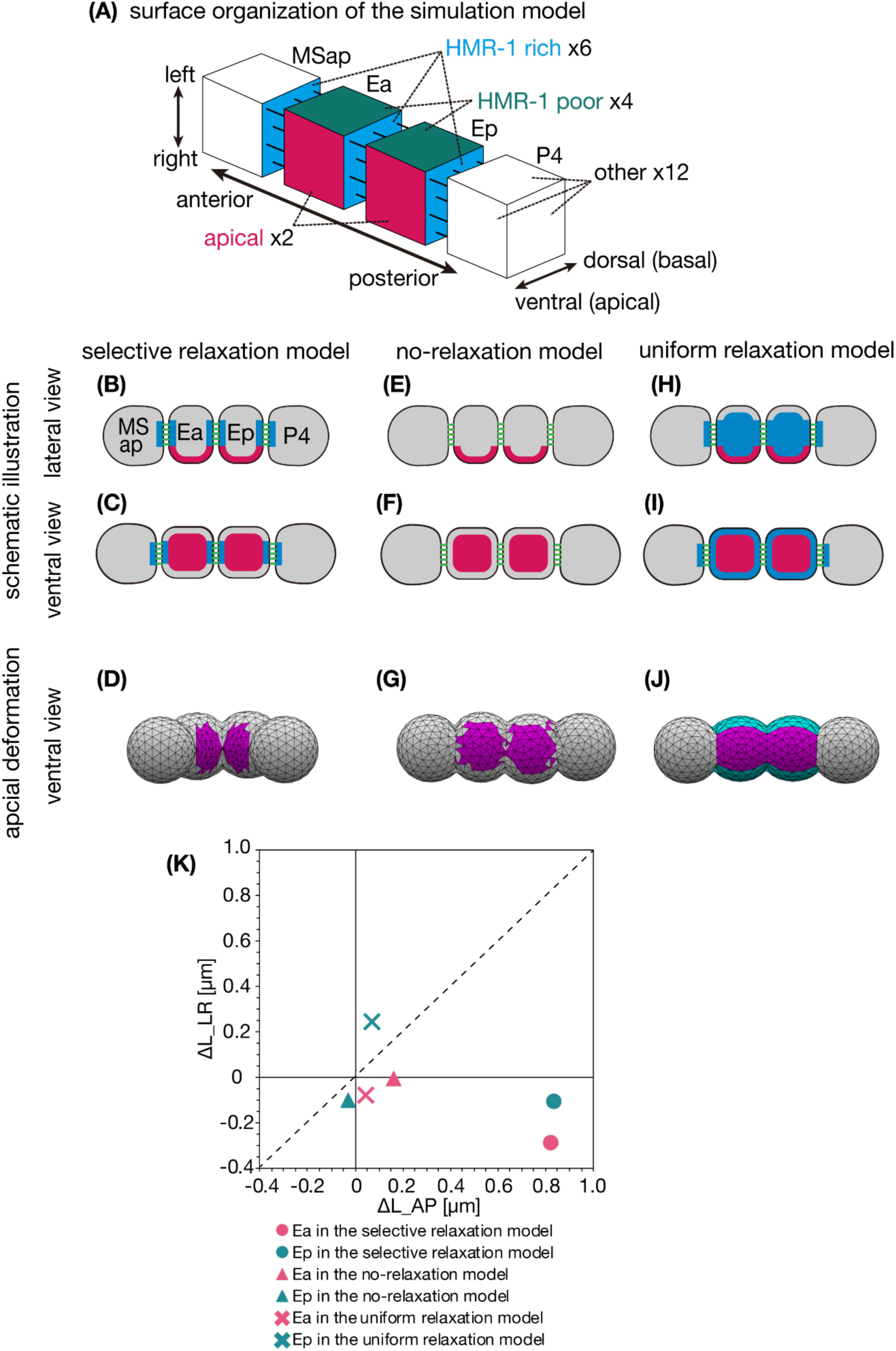
Anisotropic lateral surface tension directs anisotropic apical deformation during apical constriction. **(A)** Schematic illustration of the surface organization of the four-cell model constructed using Surface Evolver. The four cells represent MSap, Ea, Ep, and P4. Each cell–cell interface is represented by two independent contact surfaces, one from each neighboring cell. The cell surfaces are classified into four categories: apical surfaces (magenta), HMR-1-rich lateral surfaces (cyan), HMR-1-poor lateral surfaces (teal), and the remaining surfaces (“other surfaces,” white). The model comprises 24 surfaces in total: two apical surfaces, six HMR-1-rich lateral surfaces, four HMR-1-poor lateral surfaces, and 12 other surfaces. Surface tension values assigned to these surfaces in each simulation model are listed in Table 1. **(B–J)** Three mechanical models were examined: **(B–D)** the selective relaxation model, in which surface tension was reduced to 0.8 at HMR-1-rich lateral surfaces; **(E–G)** the no-relaxation model, in which no lateral surface tension was reduced; and **(H–J)** the uniform relaxation model, in which both HMR-1-rich and HMR-1-poor lateral surface tensions were uniformly reduced to 0.8. In all three models, the relative apical surface tension of Ea and Ep was progressively increased from 1.0 to 2.0. **(B, E, H)** Schematic illustration of the models viewed from the lateral side. **(C, F, I)** Corresponding schematic illustrations viewed from the ventral side. The schematics show the initial configurations before apical constriction. Surface colors indicate the assigned surface tension parameters: magenta indicates the apical surfaces of Ea and Ep, where surface tension was progressively increased from 1.0 to 2.0, cyan indicates lateral surfaces with reduced surface tension (0.8), whereas uncolored surfaces have a constant surface tension of 1.0. Green lines indicate cell–cell adhesion. **(D, G, J)** Simulated apical surface deformation viewed from the ventral side at a relative apical surface tension of 1.3. Magenta and cyan indicate the same surface tension assignments as in the schematics. **(K)** Comparison of change in apical surface dimensions along the LR and AP axes in the three simulation models as apical surface tension increased from 1.0 to 1.4. Each of the six points represents either Ea or Ep in one of the three simulation models, with the AP and LR changes plotted on the horizontal and vertical axes, respectively. Positive values indicate shortening, whereas negative values indicate elongation. Magenta and teal indicate Ea and Ep, respectively, while circles, triangles, and crosses represent the selective relaxation, no-relaxation, and uniform relaxation models, respectively. The dashed diagonal line indicates equal changes along the two axes. The selective relaxation model exhibits substantially greater shortening along the AP axis than along the LR axis, placing it below the diagonal line, whereas the no-relaxation and uniform relaxation models show little directional bias.

**Table 1.** Surface tension parameters used in the three simulation models.

|  | selective relaxation model | no-relaxation model | uniform relaxation model |
|---|---|---|---|
| *γ*<sub>apical</sub> | 1.0–2.0 | 1.0–2.0 | 1.0–2.0 |
| *γ*<sub>HMR-1-rich</sub> | 0.8 | 1.0 | 0.8 |
| *γ*<sub>HMR-1-poor</sub> | 1.0 | 1.0 | 0.8 |
| *γ*<sub>other</sub> | 1.0 | 1.0 | 1.0 |

Surface tension parameters γ<sub>apical</sub>, γ<sub>HMR-1-rich</sub>, γ<sub>HMR-1-poor</sub>, and γ<sub>other</sub> correspond to the apical surfaces, HMR-1-rich lateral surfaces, HMR-1-poor lateral surfaces, and remaining surfaces, respectively (Figure 4A). Values are expressed relative to the reference surface tension of 1.0. The apical surface tension was gradually increased from 1.0 to 2.0 in all three models.

### Video S3. Simulation of invagination in the selective relaxation model

Simulation of invagination under the surface tension conditions illustrated in Figures 4B and 4C. The relative surface tension of the apical surfaces of Ea and Ep was progressively increased from 1.0 to 2.0. Surface tension was reduced to 0.8 at three specific lateral cell–cell interfaces (MSap–Ea, Ea–Ep, and Ep–P4), while the tension of all other surfaces was maintained at 1.0. Apical surfaces are shown in magenta, lateral surfaces with reduced tension in cyan, whereas uncolored surfaces have a constant baseline tension.

### Video S4. Simulation of invagination in the no-relaxation model

Simulation of invagination under the surface tension conditions illustrated in Figures 4E and 4F. The relative surface tension of the apical surfaces of Ea and Ep was progressively increased from 1.0 to 2.0, while the tension of all other surfaces was maintained at 1.0. Apical surfaces are shown in magenta, whereas uncolored surfaces have a constant baseline tension.

### Video S5. Simulation of invagination in the uniform relaxation model

Simulation of invagination under the surface tension conditions illustrated in Figures 4H and 4I. The relative surface tension of the apical surfaces of Ea and Ep was progressively increased from 1.0 to 2.0. Surface tension was uniformly reduced to 0.8 across all lateral surfaces of Ea and Ep, including those exposed to the exterior, while the tension of all other surfaces was maintained at 1.0. Apical surfaces are shown in magenta, lateral surfaces with reduced tension in cyan, whereas uncolored surfaces have a constant baseline tension.

### Selective reduction of lateral surface tension enhances invagination driven by apical constriction

We next asked whether the selective reduction of lateral surface tension also affects the extent of invagination. In the simulations, selective relaxation of lateral surface tension not only generated anisotropic apical deformation but also markedly enhanced invagination (Figures 5A–5D and Videos S3–S5). To quantify the extent of invagination, we measured the invagination angle (θ), defined as the angle between the vector from the center of Ea to that of MSap and the vector from the center of Ep to that of P4 (Figure 5E). In the selective relaxation model, the Ea and Ep cells underwent pronounced invagination as apical surface tension increased (Figures 5B and 5F; Video S3). In contrast, the no-relaxation and uniform relaxation models showed only limited invagination (Figures 5C, 5D, and 5F; Videos S4 and S5). These results demonstrate that spatially selective regulation of lateral surface tension not only generates anisotropic apical deformation but also markedly enhances invagination driven by apical constriction, thereby linking the directionality of apical deformation to the efficiency of invagination.

**Figure 5.**
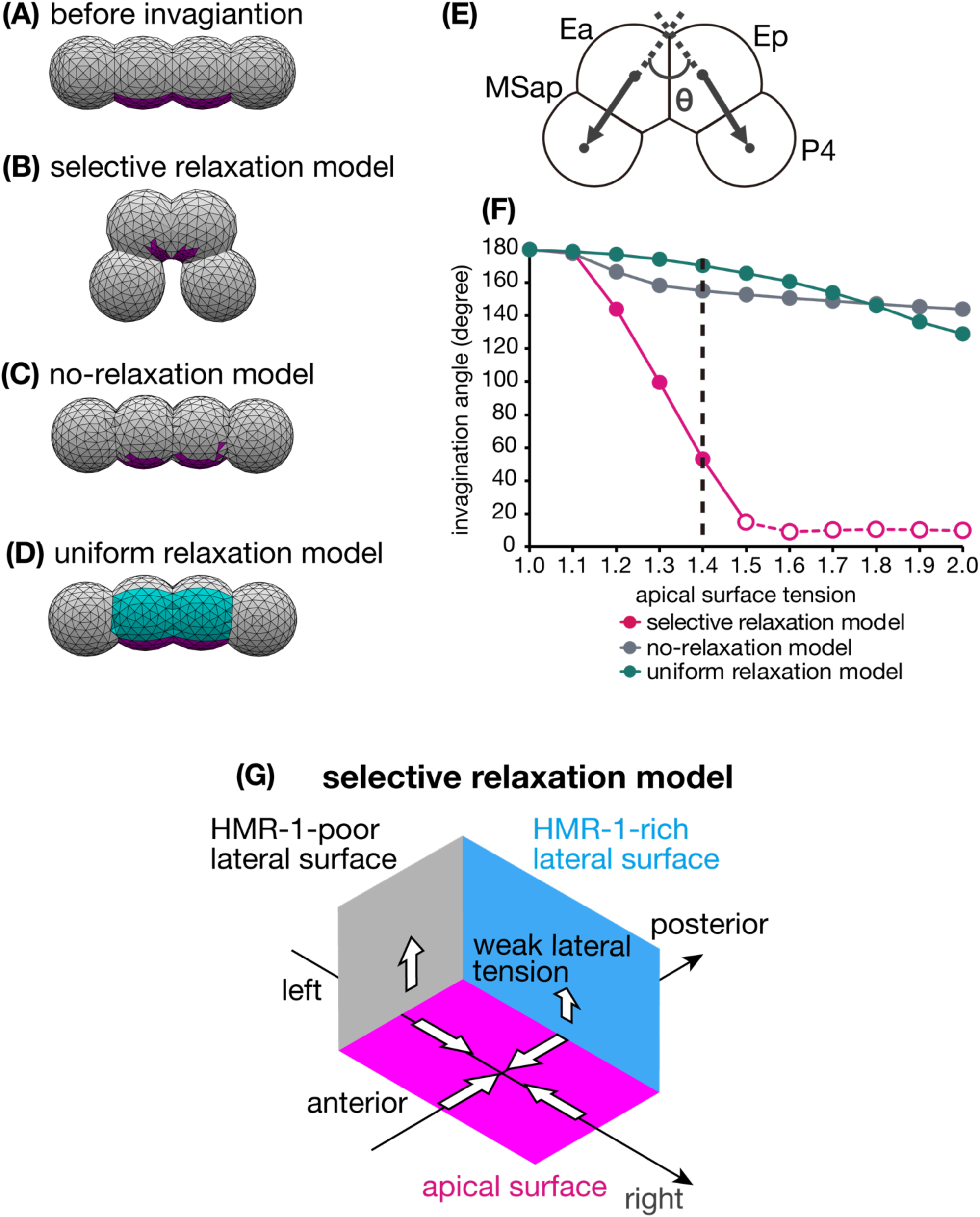
Anisotropic lateral surface tension enhances invagination driven by apical constriction. **(A)** Initial configuration of the four-cell model (MSap, Ea, Ep, and P4) before apical constriction, aligned linearly (180°). **(B–D)** Simulated configurations at a relative apical surface tension of 1.4 in the three simulation models. **(B)** The selective relaxation model produced pronounced invagination. **(C)** The no-relaxation model produced only limited invagination. **(D)** The uniform relaxation model also did not substantially enhance invagination. **(E)** Schematic illustration of the measurement of the invagination angle (θ) in Surface Evolver simulations. The angle was defined as the angle between two vectors, one directed from the center of Ea to that of MSap and the other from the center of Ep to that of P4. **(F)** Quantification of invagination as a function of relative apical surface tension in the three models. The selective relaxation model (magenta) exhibited a pronounced decrease in invagination angle as apical surface tension increased, whereas the no-relaxation model (gray) and uniform relaxation model (teal) showed only modest decreases. The vertical dashed line indicates the apical surface tension (1.4) used in (B–D). Open magenta circles indicate configurations in which Ea/Ep internalization was complete and MSap and P4 had come into contact, preventing further decreases in the invagination angle. **(G)** Schematic illustration of the proposed mechanical mechanism underlying anisotropic apical deformation and invagination. Apical constriction provides the driving force for deformation, while selective reduction of lateral surface tension at HMR-1-rich interfaces along the AP axis directs preferential shortening of the apical surfaces along the AP axis, thereby promoting invagination.

## Discussion

In this study, we show that invagination in *C. elegans* gastrulation is promoted by anisotropic regulation of lateral surface tension. By combining quantitative imaging, experimental measurements, and three-dimensional mechanical modeling, we demonstrated that anisotropic lateral surface tension generates apical deformation consistent with the anisotropic tissue deformation observed *in vivo* and markedly enhances invagination. These findings identify a previously unrecognized mechanical contribution of lateral cell–cell interfaces during epithelial folding.

### Local anisotropy as a mechanical feature of punctate invagination

A central finding of this study is that punctate invagination of the Ea and Ep cells proceeds anisotropically within the epithelial plane. Analysis of the positions of the cells surrounding Ea and Ep revealed preferential shortening along the AP axis, with much less shortening along the LR axis (Figure 1). These observations demonstrate that tissue deformation during punctate invagination exhibits a pronounced directional bias within the epithelial plane.

This finding challenges the assumption that punctate invagination can be understood as an approximately axisymmetric process around the invagination axis. Mechanical analyses of punctate invagination have traditionally focused on a single cross-sectional plane containing the apico–basal axis^13–16^, assuming that deformation outside this plane is mechanically equivalent. Our results instead demonstrate that tissue deformation within the epithelial plane is itself anisotropic and therefore cannot be fully captured by such an axisymmetric description. This distinction is particularly evident when comparing different modes of invagination. In sea urchin gastrulation, punctate invagination has generally been interpreted as an approximately axisymmetric process, whereas furrow-like invagination, exemplified by the *Drosophila* ventral furrow, is inherently directional because deformation is organized around an extended furrow^5,10,38^. Our results identify a third mechanical situation, in which punctate invagination itself proceeds anisotropically within the epithelial plane. Thus, in addition to determining whether epithelial tissues bend, morphogenetic mechanisms may also regulate the direction in which punctate invagination preferentially proceeds. Anisotropy should therefore be considered an additional mechanical degree of freedom in epithelial folding. Rather than representing a simple deviation from symmetry, anisotropy can be an important mechanical feature of punctate invagination that can influence both the direction and the efficiency of tissue folding.

### Apical constriction generates contractile force, whereas lateral mechanics determine deformation

Our results also refine the current mechanical interpretation of apical constriction. Although neighboring cells converged preferentially along the AP axis, laser ablation revealed no detectable anisotropy in apical cortical recoil. Thus, the direction of apical deformation cannot be explained simply by anisotropic actomyosin contractility.

Apical actomyosin contraction generates forces that are transmitted through cell–cell junctions to neighboring cells, contributing to tissue deformation^21,23^. Our simulations indicate that directional deformation emerges from the mechanical environment surrounding the apical surfaces. When lateral surface tension was reduced selectively at AP-oriented interfaces, the apical surfaces became shortened along the AP axis, consistent with the preferential AP convergence observed *in vivo* (Figures 4D and 4K). In contrast, uniform lateral surface tension failed to generate comparable deformation or enhance invagination (Figures 4G, 4J, 5C and 5D). These findings suggest that apical actomyosin contractility provides the driving force for constriction, whereas the spatial distribution of lateral surface tension determines how this force is translated into directional tissue deformation. Thus, apical contractility and lateral interfacial mechanics act cooperatively to generate three-dimensional tissue deformation.

### A second mechanical function of E-cadherin during gastrulation

Previous studies established that E-cadherin-mediated adhesion is essential for *C. elegans* gastrulation^22^ and proposed that E-cadherin promotes invagination by mechanically coupling the apical actomyosin network to cell–cell junctions^21^. In this molecular clutch model, the coupling enables contractile forces generated in Ea and Ep to be transmitted to neighboring cells. These forces can drive the movement of neighboring cells, contributing to closure of the opening at the embryo surface as Ea and Ep internalize, without necessarily requiring active contraction by the neighboring cells themselves.

Our findings suggest an additional mechanical role for E-cadherin at lateral cell–cell interfaces. E-cadherin-mediated adhesion can stabilize and expand cell–cell contact surfaces, thereby reducing interfacial surface tension. We found that HMR-1 is preferentially enriched at specific lateral interfaces, where junctional geometry was consistent with reduced interfacial surface tension *in vivo*. Incorporating this experimentally observed anisotropy into our mechanical model was sufficient to reproduce both enhanced invagination and the anisotropic deformation of apical surfaces. Thus, the molecular clutch model and the mechanism proposed here describe complementary aspects of E-cadherin’s role during gastrulation. Whereas the clutch model explains how apical contractile forces are mechanically coupled to cell boundaries and transmitted to neighboring cells, our model suggests how the spatial regulation of interfacial surface tension biases the resulting tissue deformation. Together, these results extend the current view of E-cadherin’s role by demonstrating that cell adhesion contributes not only to mechanical coupling but also to the spatial organization of tissue mechanics.

### Spatial organization of mechanics as a general principle of morphogenesis

An important implication of this study is that the direction of tissue deformation during apical constriction can be influenced by the spatial distribution of lateral interfacial surface tension. Previous studies have shown that anisotropic epithelial deformation can arise from planar-polarized actomyosin contractility^39^ or mechanical constraints imposed by tissue-level tension^11^. Studies of adhering vesicle aggregates have also shown that differences in interfacial surface tension can influence the morphology of connected structures^40^. Our results identify spatial differences in lateral interfacial surface tension as an additional mechanical factor contributing to anisotropic tissue deformation.

Although our study focuses on *C. elegans* gastrulation, spatial patterning of interfacial mechanics may represent a general mechanism for directing epithelial morphogenesis. Many morphogenetic events involve tissues with intrinsic planar polarity or regional differences in cell adhesion. Our results suggest that local modulation of lateral interfacial mechanics may direct tissue deformation without requiring corresponding anisotropy in force generation itself.

Our computational framework also provides a useful strategy for investigating such questions. Unlike conventional shared-surface models, our model represents the surfaces of neighboring cells independently and connects them through elastic elements, allowing mechanical coupling and surface tension to be controlled separately. This approach enables us to examine the contribution of spatially patterned lateral surface tension while maintaining mechanical connectivity between cells. Similar approaches may prove useful for dissecting the contributions of cell adhesion and interfacial mechanics in other morphogenetic systems.

### Limitations and future perspectives

Several limitations should be noted. First, interfacial surface tension was inferred indirectly from junction geometry rather than measured directly. Future approaches capable of directly quantifying interfacial tension will further test the proposed mechanism. Second, although the preferential enrichment of HMR-1 at specific lateral cell–cell interfaces is central to our proposed mechanism, how this spatial distribution of HMR-1 is established remains unknown. Third, our mechanical model incorporates specific constraints on cell geometry and connectivity. Although these constraints were identical across the simulation models, it remains unclear whether the anisotropic deformation generated by selective relaxation would persist under alternative constraints, such as conservation of cell surface area or different restrictions on vertex displacement. Future simulations examining these conditions will help assess the generality of the proposed mechanism. In summary, our study identifies anisotropic regulation of lateral surface tension as a previously unrecognized mechanism that promotes epithelial invagination. More broadly, these findings suggest that tissue morphogenesis is governed not only by the generation of contractile forces but also by the spatial organization of the mechanical environment through which those forces are translated into directional tissue deformation.

## Materials and Methods

### Establishment and maintenance of *Caenorhabditis elegans* strains

The *C. elegans* strains used in this study are listed in Table S1. Worms were maintained at 22 °C on nematode growth medium (NGM) agar plates seeded with *Escherichia coli* OP50 according to standard procedures^41^. The CAL3031 strain was generated in this study using CRISPR/Cas9-mediated genome editing following a previously reported procedure^42^. A guide RNA targeting the C-terminal region of *nmy-2* (5′-GTTAGTTGCGAACTGAGTCG-3′) was cloned into the pJW1219 vector. The modified pJW1219 plasmid and repair templates containing mBaoJin^43^ and *nmy-2* sequences, generated using pDFT vectors, were co-injected into the gonads of young adult N2 hermaphrodites together with pRF4, a co-injection marker used to identify progeny carrying injected DNA based on the Roller phenotype. Following screening for the Roller phenotype, animals were further screened for fluorescence. The CAL3041 strain was generated by crossing CAL3031 with ZZY0637^44^. F3 animals homozygous for both *wj222* and *zzyIs139* were identified by confirming that all their F4 progeny were positive for both NMY-2::mBaoJin and PH::mCherry fluorescence. The CAL3081 strain was generated by crossing LP172^45^ and ZZY0637. F2 animals homozygous for both *cp21* and *zzyIs139* were identified by confirming that all their F3 progeny were positive for both HMR-1::GFP and PH::mCherry fluorescence.

**Table S1.**
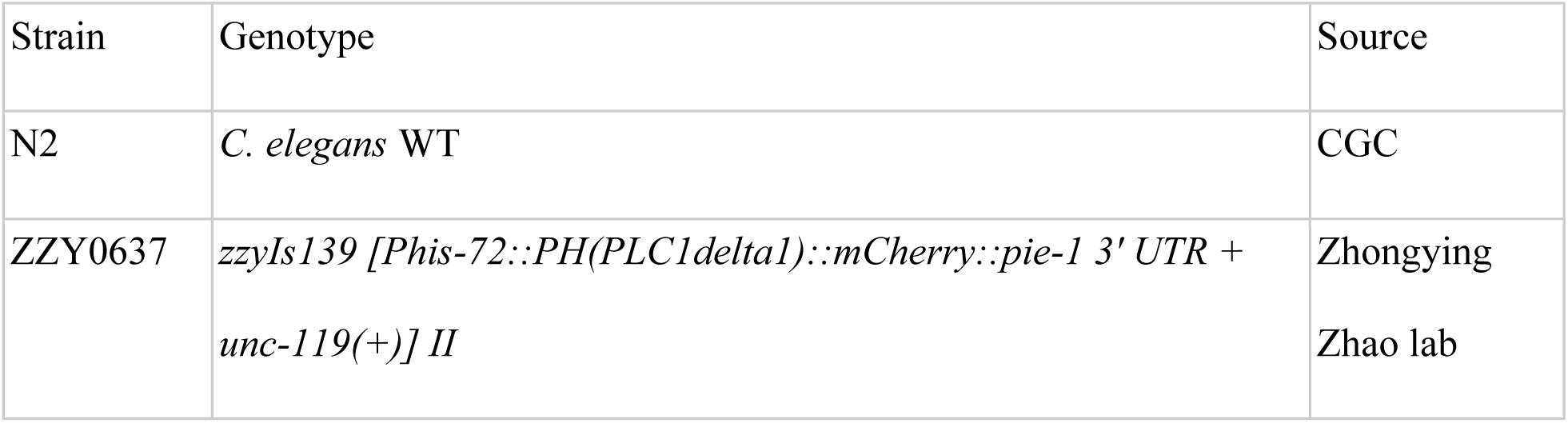
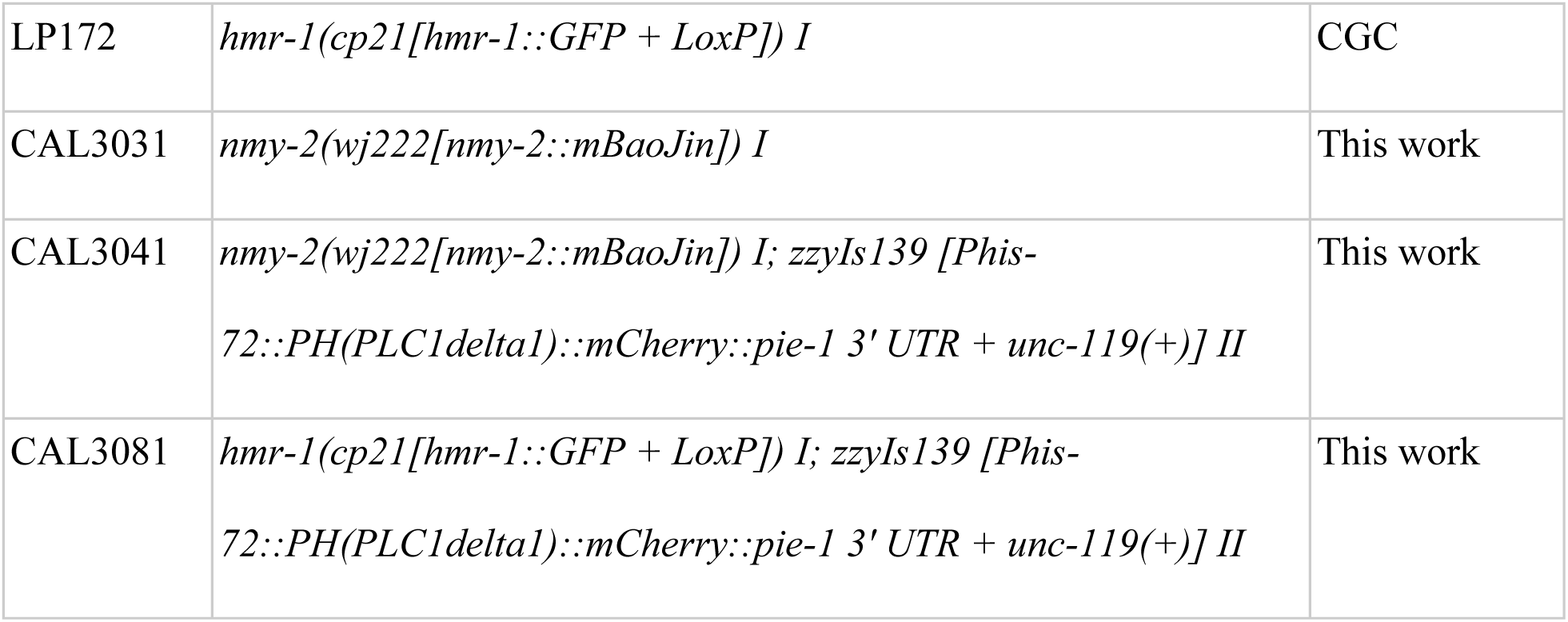
The *C. elegans* strains used in this study.

| Strain | Genotype | Source |
|---|---|---|
| N2 | *C. elegans* WT | CGC |
| ZZY0637 | *zzyIs139 [Phis-72::PH(PLC1delta1)::mCherry::pie-1 3' UTR + unc-119(+)] II* | Zhongying Zhao lab |
| LP172 | *hmr-1(cp21[hmr-1::GFP + LoxP]) I* | CGC |
| CAL3031 | *nmy-2(wj222[nmy-2::mBaoJin]) I* | This work |
| CAL3041 | *nmy-2(wj222[nmy-2::mBaoJin]) I; zzyIs139 [Phis-72::PH(PLC1delta1)::mCherry::pie-1 3' UTR + unc-119(+)] II* | This work |
| CAL3081 | *hmr-1(cp21[hmr-1::GFP + LoxP]) I; zzyIs139 [Phis-72::PH(PLC1delta1)::mCherry::pie-1 3' UTR + unc-119(+)] II* | This work |

### Preparation and mounting of the embryos for live cell imaging

For live imaging, adult hermaphrodite worms were dissected in 0.75× egg salt, and embryos were collected. For lateral-view imaging, four-cell-stage embryos were placed on a 2.5% agarose pad with 0.75× egg salt and covered with an 18×18 mm coverslip (Matsunami glass). To visualize embryos from a ventral view, a 5.2 µL drop of 0.75× egg salt was placed onto a poly L-lysine–coated 18×18 mm coverslip (Matsunami Glass). Four-cell-stage embryos were placed on the glass and manually oriented with their ventral side facing the coverslip. The embryos were allowed to develop to the eight-cell stage in a humidity chamber. They were then gently covered with a coverslip (Matsunami Glass) and sealed with VALAP (a 1:1:1 mixture of Vaseline [petroleum jelly], lanolin, and paraffin). To examine junctional geometry (Figure 3C), eggshell was removed from a two-cell-stage embryo using previously reported methods^46^. Following eggshell removal, the AB and P1 blastomeres were separated by repeated pipetting through a micropipette with an approximately 30-μm inner diameter, prepared by pulling a glass capillary (GD-1; Narishige, Tokyo, Japan) using a micropipette puller (P-1000IVF; Sutter Instrument, Novato, CA). The isolated P1 blastomere was cultured to generate descendant cells, including MSxx, Ea, Ep, and P4.

### Live cell imaging, without laser ablation

Embryos were imaged at approximately 22 °C using a spinning-disk confocal microscope unit (CSU-MP^47^; Yokogawa Electric) mounted on an IX71 microscope (Olympus) equipped with a 40× objective lens and an EM-CCD camera (iXon; Andor). Imaging acquisition was controlled using NIS-Elements software (Nikon).

### Laser ablation experiments

Embryos were observed using a spinning-disk confocal microscope (CSU-W1; Yokogawa Electric) equipped with a CMOS camera (ORCA-Fusion BT, Hamamatsu photonics), a 100× UPLAPO OHR oil-immersion objective (N.A. = 1.50; Evident), LDI Prime laser-diode system (Oxxius) comprising 488 nm (1 W) and 555 nm (1 W) lasers, mounted on a Ti2-E microscope (Nikon) and controlled using MetaMorph at approximately 22 °C. Embryos were mounted in the ventral orientation, and laser irradiation was performed 22 min after Ea/Ep birth, corresponding to the time of the most rapid decrease in apical surface area on average across embryos. A point-ablation assay was performed at the center of the apical surface of the Ep cell with 30% laser power, 1 thickness, and 20 repetitions using a 355-nm UV pulse laser (2 kW, 1 μJ energy, 400 ps, 20 kHz, Teem Photonics). The laser irradiation site was set as a square with a length of 0.34 μm at the center of the apical surface of the Ep cell. Images were acquired at 1-s intervals from 10 s before to 15 s after the irradiation.

### Quantification

#### (a) Cell movements during gastrulation (Figure 1)

Images were acquired using a 561 nm laser with 144-ms exposure using the setup of the live-cell imaging procedure. Z-stack images were taken at 0.5-μm intervals on the z-axis. Time-lapse images were collected at 5 min intervals for less than 1 h. Under these microscopic conditions, *C. elegans* embryos were confirmed to hatch. Individual cells from PH::mCherry (strain ZZY0637) fluorescence images were manually segmented, and the three-dimensional center-of-mass coordinates (X, Y, Z) for each cell were obtained in Imaris (Bitplane). We analyzed the six cells surrounding Ea and Ep and calculated the intercellular distances for all 15 cell pairs from their three-dimensional center-of-mass coordinates. The intercellular distances for each cell pair were separated into the AP and LR axes, and the intercellular distances in each direction were calculated. These values were determined for the initiation and completion of Ea/Ep invagination, respectively. Changes in intercellular distances were calculated by subtracting the distance at the completion of Ea/Ep invagination from that at its initiation, yielding ΔD_AP and ΔD_LR for the AP and LR directions, respectively. Positive and negative values indicate decreases and increases in intercellular distance, respectively.

#### (b) HMR-1 localization (Figure 3B)

Images were acquired using a 480 nm laser with 96-ms exposure and a 561 nm laser with 48-ms exposure using the setup of the live cell imaging procedure. Z-stack images were taken at 0.5-μm intervals on the z-axis. Time-lapse images were collected at 1 min intervals for less than 1 h. Under these microscopic conditions, *C. elegans* embryos were confirmed to hatch. HMR-1::GFP fluorescence at cell–cell contact surfaces was quantified from ventral view images (strain CAL3081). Fluorescence intensity was measured using ImageJ/Fiji by drawing linear ROIs along individual cell–cell contact surfaces and manually subtracting the average gray value of the background region. The fluorescence intensity was averaged over time from the birth of the MS granddaughter cell to the division of Ea and Ep.

Statistical significance was assessed using prespecified one-sided exact permutation tests based on within-embryo comparisons (n = 5 embryos). For Ea-associated boundaries, the three Ea–MS boundaries (Ea–MSap, Ea–MSpa, and Ea–MSpp) were compared pairwise with the two Ea–AB boundaries (Ea–ABplpa and Ea–ABplpp), yielding six comparisons per embryo. For Ep-associated boundaries, Ep–P4 was compared with Ep–ABplpp and Ep–Cap, yielding two comparisons per embryo. The total number of comparisons in which the Ea–MS or Ep–P4 boundary had the higher fluorescence intensity was used as the test statistic. Exact null distributions were obtained by exhaustively permuting the boundary labels within each embryo. For the Ea–Ep analysis, its fluorescence intensity was ranked among the nine cell–cell boundaries in each embryo, with rank 1 assigned to the highest intensity, and the ranks were summed across the five embryos. The exact null distribution of the rank sum was obtained by exhaustively permuting the Ea–Ep label within each embryo. The Ep non-contact surface, Ep–(–), was shown for reference but was excluded from all statistical comparisons. P < 0.05 was considered statistically significant.

#### (c) Junction angles (Figure 3C)

To assess differences in surface tension, we measured the junction angles at the Ea–Ep interface (Φ₁) and at the interface between P4 and its neighboring cell (Φ₂). For each angle, a circle with a radius of 3 µm was centered at the vertex where the cell boundaries meet. The two points at which the circle intersected the cell boundaries defining the angle were identified, and the angle between the lines connecting these points to the vertex was measured.

#### (d) Directional bias of NMY-2 recoil (Figure 2)

Cortical recoil following laser ablation was quantified by NMY-2 signal patterns (strain CAL3041). To visualize cortical NMY-2 recoil, time-lapse images were color-coded according to acquisition time and overlaid into a single image. To quantify the recoil direction, displacement vectors were generated from the NMY-2 signal patterns before and after each time interval. The displacement vectors were expressed in a coordinate system centered on the ablation site, with the AP axis defined as the x-axis and the LR axis as the y-axis. The displacement vectors were reflected across the x-axis and y-axis to map them into the first quadrant. The mean displacement vector was then calculated from all transformed vectors. The angle between the mean displacement vector and the x-axis indicates the preferred recoil direction. Angles between 0° and 90° indicate net movement away from the ablation site, with 0° corresponding to the AP axis, 90° to the LR axis, and 45° indicating no directional bias between the two axes. Since the magnitude of the mean displacement vector reflects the overall degree of cortical recoil, we used the angle at which the magnitude of the mean displacement vector was greatest during the 3 seconds following laser ablation as an indicator to assess the directional bias of recoil.

### Mechanical modeling using Surface Evolver

#### (a) Construction of the four-cell system in *C. elegans* gastrulation

To investigate the effects of surface tension at specific contacts on invagination during *C. elegans* gastrulation, simulations were performed using Surface Evolver version 2.70^31,35^. Surface Evolver computes equilibrium shapes of triangulated surfaces by minimizing the total mechanical energy under specified geometrical constraints.

Our four-cell system consists of four cells: MSap, Ea, Ep, and P4 (Figure 4A). Each cell–cell interface is represented by two independent contact surfaces, one from each neighboring cell, connected with elastic elements. The 24 cell surfaces were classified into four categories: apical surfaces of Ea and Ep (magenta in Figure 4A), HMR-1-rich lateral surfaces (cyan), HMR-1-poor lateral surfaces (teal), and the remaining (“other surfaces”, white).

The dimensional mechanical energy is a sum of apical, lateral including HMR-1-rich and HMR-1-poor surfaces, and other surfaces, as indicated below:

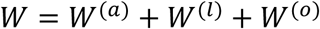

The energy at the apical surfaces reads,

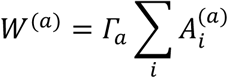

Here, *Γ_a_* denotes the surface tension at the apical side, 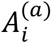 denotes the apical area of cell *i*. The energy at the lateral surfaces reads,

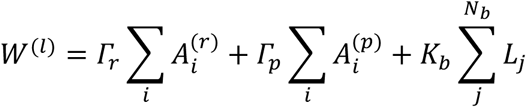

Here, *Γ_r_* denotes the surface tension of the HMR-1-rich lateral surfaces, and 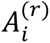 denotes its surface area. In the second term, *Γ_p_* denotes the surface tension of the HMR-1-poor lateral surfaces, and 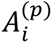 denotes its surface area. In the third term, *K_b_* represents the line tension of elastic elements connecting between the two adjacent HMR-1-rich lateral surfaces, thus penalizing the length of elastic elements *L_j_*. These elastic elements are initially created by connecting the vertices having the same position. After the mesh refining, the total number of bonds amounts to *N_b_*.

The energy of other surface reads,

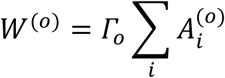

Here, *Γ_o_* denotes the surface tension of other surfaces, and 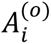 denotes its surface area. Cell volume was constrained to *V_o_* = 1.0 during simulation. We define the dimensionless energy as 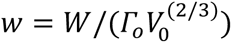 that reads,

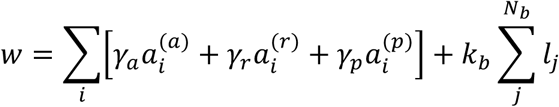

Here the dimensionless surface tension *γ_a_*, *γ_r_*, and *γ_p_* are defined as *Γ_a_*/*Γ_o_*, *Γ_r_*/*Γ_o_*, and *Γ_p_*/*Γ_o_*, respectively. The dimensionless surface areas 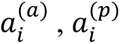, and 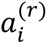 are defined as 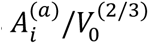, 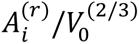, and 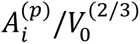, respectively. Similarly, the dimensionless line tension *k_b_* and bond length *l_j_* are defined as 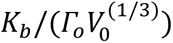 and 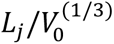, respectively. In the simulation, apical surface tension was gradually increased from 1.0 to 2.0. For each tension value, the total mechanical energy was minimized using Surface Evolver over 2,000 iterations, resulting in a total of 22,000 iterations for each simulation condition. Vertex averaging and mesh equiangulation were applied during minimization to obtain a quasistatic equilibrium shape. We examined three simulation conditions, with the corresponding parameters listed in Table 1.

#### (b) Two cell models for characterizing cell–cell adhesion

To examine whether independently represented contact surfaces connected by elastic elements reproduce the mechanical behavior of a shared cell–cell interface (Figure S2), we constructed two two-cell models using Surface Evolver. In the first model, the two cells shared a single interface. In the second model, each cell had an independent contact surface, and the two surfaces were connected by short elastic elements. The surface tension of the contact surfaces was set to 0.8, 1.0, or 1.2, while that of non-contact surfaces was maintained at 1.0. Equilibrium configurations were quantified using the θ/2 method, in which the contact angle is estimated from the height and base radius of an approximately spherical cell or droplet^48^.

#### (c) Changing surface tension at apical surface and lateral surfaces

Apical constriction was simulated by gradually increasing the surface tension of the Ea and Ep apical surfaces from 1.0 to 2.0. To evaluate the effects of the spatial distribution of lateral surface tension, we examined three simulation models (Table 1). In the no-relaxation model, all non-apical surface tensions were maintained at 1.0 (Table 1). In the selective relaxation model, the surface tension of the six HMR-1-rich surfaces was reduced to 0.8, while all remaining non-apical surface tensions were maintained at 1.0 (Table 1). In the uniform relaxation model, the surface tension of both HMR-1-rich and HMR-1-poor lateral surfaces was reduced to 0.8, while those of other surfaces were maintained at 1.0 (Table 1).

#### (d) Quantification of directional changes in apical surface dimensions in simulations (Figure 4K)

Directional changes in the apical surface dimensions of Ea and Ep were quantified in the three simulation models (selective relaxation, no-relaxation, and uniform relaxation). The coordinates of all vertices constituting each apical surface were extracted from the simulated cell meshes. The AP and LR dimensions were defined as the differences between the maximum and minimum vertex coordinates along the AP-and LR axes, respectively. These dimensions were measured at the initial (γ_apical_ = 1.0) and final (γ_apical_ = 1.4) states. Directional shortening was calculated by subtracting the final dimension from the initial dimension along each axis, yielding ΔL_AP and ΔL_LR. Positive values indicate shortening, whereas negative values indicate elongation.

#### (e) Invagination angle in Surface Evolver simulations (Figure 5E)

The extent of invagination in Surface Evolver simulations was quantified by measuring the invagination angle (θ). The center of each cell (MSap, Ea, Ep, and P4) was defined as the mean position of all vertices constituting that cell. The invagination angle was calculated as the angle between two vectors: one pointing from the center of Ea to that of MSap, and the other from the center of Ep to that of P4.

## Declaration of generative AI and AI-assisted technologies in the manuscript preparation process

During the preparation of this work, the authors used ChatGPT (OpenAI) to assist with generating Python code for data analysis, translating portions of the manuscript into English, and improving the language and readability of the manuscript. After using this tool, the authors reviewed and edited the content as needed and take full responsibility for the content of the published article.

## Author Contributions

Conceptualization: T.K. and A.K.

Methodology: T.K., K.Y., T.C., M.I., F.M., and A.K.

Software: T.K. and T.C.

Investigation: T.K.

Formal analysis: T.K.

Resources: T.K., K.Y., T.C., M.I., F.M., and A.K.

Visualization: T.K.

Writing – original draft: T.K. and A.K.

Writing – review & editing: T.K., K.Y., T.C., M.I., F.M., and A.K.

Supervision: M.I., F.M., and A.K.

Funding acquisition: T.K. and A.K.

## Acknowledgements

We thank Dr. Takayuki Torisawa for his advice on constructing the mechanical model and his assistance in experiments to establish the CAL3031 strain; Drs. Kenji Kimura and Yukako Nishimura for the advice in laser ablation experiments; and members of the Cell Architecture Laboratory and the Physics and Cell Biology Laboratory in the National Institute of Genetics, as well as members of the Motegi Laboratory at Hokkaido University, for their valuable feedback on this study. We also thank Dr. Zhongying Zhao for providing the ZZY0637 strain. The LP172 strain was provided by the *Caenorhabditis* Genetics Center (CGC), which is funded by NIH Office of Research Infrastructure Programs (P40 OD010440). This work was supported by JST SPRING Grant Number JPMJSP2104 for TK, by JSPS KAKENHI Grant Numbers 16H00816, 25K02276 for AK, and by the Grant for Joint Research Program of Institute for Genetic Medicine, Hokkaido University.

